# Decoupling Structure and Property in Discrete Protein Diffusion: Alignment Dynamics and Collapse Mechanisms

**DOI:** 10.64898/2026.02.05.701726

**Authors:** Junqi Wu, Liyan Dong, Nuo Jia, Longyi Li, Hao Zhang

## Abstract

Direct Preference Optimization (DPO) has emerged as a powerful paradigm for aligning generative models, yet its *temporal optimization dynamics* in the discrete diffusion space of proteins remain poorly understood. Existing approaches often assume that maintaining structural integrity while optimizing physicochemical properties requires simultaneous, tightly coupled reinforcement learning constraints. In this work, we challenge this assumption by uncovering a fundamental *temporal decoupling* between structural and functional alignment. Using antibody design as a testbed, extensive trajectory analysis reveals two distinct regimes: (1) *Instant Structural Alignment*, where the strong generative prior of discrete diffusion rapidly eliminates structural hallucinations via denoising within the first few epochs; and (2) *Slow Property Adaptation*, where physicochemical attributes improve gradually over a prolonged optimization window. We further identify a critical transition point around Epoch 50, which empirically defines a Pareto-optimal boundary between property improvement and structural stability. Beyond this point, continued optimization induces a sharp phase transition into a *Structural Collapse* regime. To isolate the physical driver underlying this collapse, we introduce a counterfactual preference experiment targeting negative charge. We observe a striking *symmetrical collapse*: while hydrophilicity optimization induces a Poly-Arginine (+) degeneration, negative charge optimization drives a Poly-Aspartate (-) degeneration. Despite opposite physicochemical trajectories, including extreme shifts in isoelectric point (*>* 11 vs. *<* 4.5), both regimes converge to the same structural failure. This symmetry demonstrates that generic Coulombic repulsion, rather than residue-specific bias, constitutes the fundamental physical constraint being violated. Our findings reveal that discrete diffusion models possess strong intrinsic structural robustness, enabling minimalist alignment strategies provided optimization halts before this physical boundary. More broadly, this work offers a mechanistic warning against unchecked reward optimization in biological generation, illustrating a concrete manifestation of Goodhart’s Law in protein design. **Code and data are available at** https://github.com/Wu-Junqi/DPO-Protein-Diffusion.

## 1 Introduction

Generative protein design promises to transform molecular engineering [1] by producing sequences that satisfy both rigorous structural constraints [2] and functional objectives. A prevalent assumption in current literature is that structural and functional alignments represent equally challenging optimization hurdles. Consequently, methods often resort to elaborate strategies—such as physics-informed rewards [3] or auxiliary losses [4]—which increase complexity and obscure the intrinsic behavior of the generative model. For instance, while continuous diffusion models like RFdiffusion [5] excel in backbone generation, discrete variants [6] remain underexplored in terms of their alignment dynamics, leaving significant gaps in understanding how generative priors interact with property objectives over time.

In this work, we challenge this complexity assumption by advocating for a return to minimalism. We posit that the generative priors of discrete diffusion models possess an underestimated structural robustness, potentially enabling rapid structural alignment without auxiliary engineering. **Our empirical baseline supports this hypothesis, suggesting that the prior alone acts as a potent structural denoiser capable of rapid manifold projection**.

This observation leads to our central thesis: *Structural alignment is not the bottleneck; the true challenge lies in the dynamic conflict between structure and property*. This decoupling is particularly relevant in the era of advanced structure prediction tools like AlphaFold [7], which have revolutionized our ability to evaluate generated sequences against realistic structural constraints. Despite advances, existing studies often evaluate alignment at convergence without dissecting temporal trajectories, implicitly assuming synchronized improvement. This oversight raises critical questions: How do structural priors and property objectives evolve on distinct timescales in discrete diffusion settings? What microscopic mechanisms drive potential instabilities when these objectives conflict?

To address this gap, we analyze alignment trajectories by subjecting the model to property-focused Direct Preference Optimization (DPO) [8], investigating a potential **Dynamics Decoupling** phenomenon (schematically illustrated in Figure 1(B)). Rather than assuming coupled processes, our trajectory-level analysis explores whether alignment manifests through orthogonal timescales: an initial restoration phase driven by the prior, a transient Pareto-optimal window, and a final phase susceptible to reward hacking and chemical degeneration.

**Figure 1.**
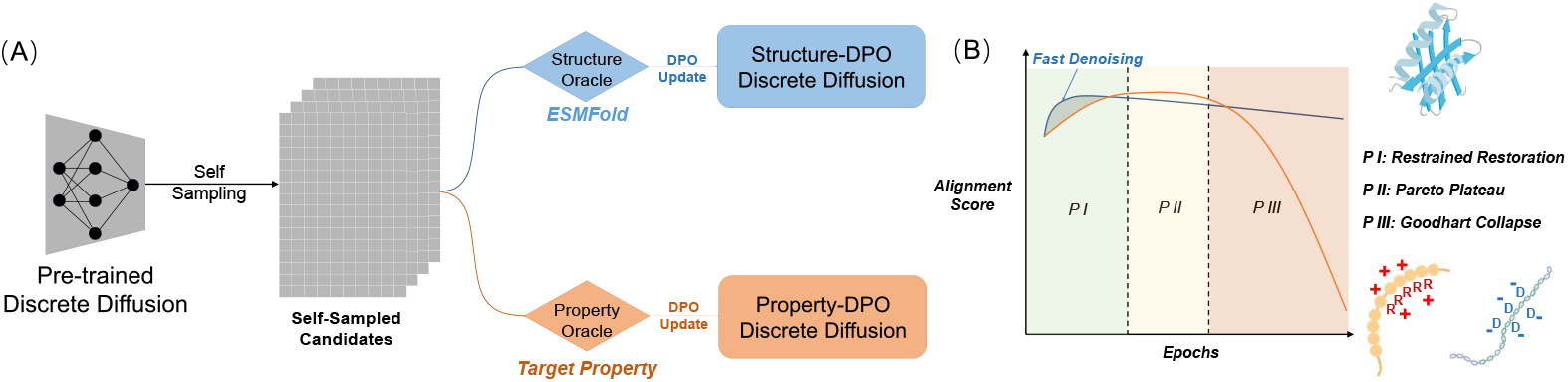
The Minimalist Alignment Protocol and Dynamics Decoupling. (A) Pipeline using discrete diffusion and deterministic oracles (Target Property). (B) Schematic of Alignment Dynamics. The model evolves through Restrained Restoration (I) and a Pareto Plateau (II) before succumbing to Goodhart Collapse (III). **Note:** As illustrated in the bottom-right, the collapse manifests symmetrically: optimizing hydrophilicity drives **Poly-Arginine (+)** accumulation, while optimizing negative charge drives **Poly-Aspartate (-)** accumulation, both triggering electrostatic repulsion.

To rigorously investigate these dynamics, we adhere to a strictly minimalist protocol: standard discrete diffusion backbone, offline DPO, and deterministic oracles. Our contributions are threefold:

- **Uncovering Dynamics Decoupling:** We empirically explore how structural restoration and property adaptation operate on distinct timescales, challenging the prevailing need for complex coupled objectives.
- **Mechanism Verification via Symmetrical Collapse:** By introducing a counterfactual *Negative Charge* experiment, we reveal a symmetrical failure mode (Poly-Arginine vs. Poly-Aspartate). This confirms that generic Coulombic repulsion, rather than specific residue bias, is the fundamental physical constraint.
- **A Robust Minimalist Baseline:** We establish that standard DPO, stabilized by a synchronized masking scheme, offers competitive performance without intricate RL, serving as a strong foundation for future minimalist pipelines.

## 2 Related Work

### 2.1 Generative Protein Design with Diffusion

Deep generative models have fundamentally transformed protein engineering. Early approaches relied on autoregressive language models or GANs, but diffusion models have recently emerged as the dominant paradigm due to their stable training objectives and high-fidelity generation.

#### Continuous vs. Discrete Diffusion

In the continuous domain, methods operating on SE(3) manifolds, such as RFdiffusion [5], Chroma [9], and FrameDiff [10], have achieved breakthrough performance in backbone generation. These models excel at capturing geometric constraints but often require a separate inverse folding step (e.g., ProteinMPNN) to recover sequences. Conversely, discrete diffusion frameworks [6, 11] operate directly in the amino acid sequence space. Models like EvoDiff [12], HuAbDiffusion [13], and MDLM [14] formulate generation as an iterative denoising process over masked tokens. By leveraging the discrete nature of sequences, they provide a strong syntactic prior akin to Masked Language Models (MLMs) [15]. However, while these works focus heavily on maximizing generation fidelity (e.g., perplexity or recovery rate), the *temporal dynamics* of how these priors interact with downstream alignment objectives remain largely unexplored.

### 2.2 Preference Alignment in Biology

While pre-trained models capture the distribution of natural proteins, engineering functional proteins requires aligning these models with specific physicochemical properties (e.g., stability, binding affinity).

#### From RLHF to DPO

Alignment techniques have evolved from Reinforcement Learning from Human Feedback (RLHF) [16], which typically employs Proximal Policy Optimization (PPO) [17]. However, PPO is notoriously unstable and computationally expensive due to the need for value networks and online rollouts. Direct Preference Optimization (DPO) [8] has recently emerged as a compelling alternative, optimizing the policy directly from preference data without explicit reward modeling.

#### Applications and Risks

In the biological domain, these methods are increasingly applied to optimize molecular properties. Recent works like Physio-DPO [3] and PocketX [18] demonstrate the efficacy of preference learning in structure-based design. However, forceful optimization against surrogate reward models (oracles) carries significant risks. As noted in the broader AI alignment literature, unchecked optimization leads to *Goodhart’s Law* [19] and *reward hacking* [20], where the model exploits adversarial loopholes in the reward function. In protein design, this often manifests as the generation of sequences that are numerically optimal but biophysically invalid (e.g., unfolded or chemically extreme).

### 2.3 Dynamics of Generative Optimization

Understanding the training dynamics of generative models provides crucial insights into their failure modes. In computer vision, empirical studies have revealed a “coarse-to-fine” generation hierarchy in diffusion models [21, 22], where global structure is determined early, followed by local texture refinement. In contrast, the alignment dynamics of discrete protein diffusion remain opaque. Existing studies [23] typically report only final converged metrics, obscuring the “tug-of-war” between the generative prior and the alignment objective. Our work bridges this gap by systematically dissecting the temporal trajectories of alignment, revealing the critical decoupling that governs structural stability.

## 3 A Minimalist Protocol for Alignment Dynamics

To investigate the intrinsic alignment behavior of discrete diffusion models, we deliberately adopt a minimalist experimental protocol, as depicted in Figure 1(A). Rather than proposing intricate architectures or auxiliary dense reward models, our goal is to isolate and observe alignment dynamics under the weakest possible intervention. This section details the protocol, designed as a controlled observation framework rather than an algorithmic contribution.

### 3.1 Preliminaries: Discrete Diffusion Priors

We consider generative modeling over protein sequences defined on a discrete state space 𝒳 = 𝒱^*L*^, where 𝒱 denotes the amino acid vocabulary augmented with a special `[MASK]` token.

#### Forward Process

The forward diffusion process progressively corrupts a clean sequence *x*_0_ into noise via an absorbing state formulation [6]. The forward process is implemented via stochastic masking, which corresponds to the absorbing-state transition defined in Eq. (1). This formulation treats generation as iterative parallel decoding, conceptually similar to MaskGIT [24] and EvoDiff [12].

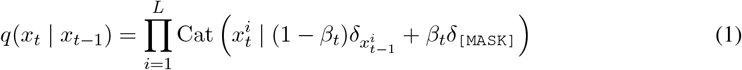

where *β*_*t*_ is the schedule controlling the masking rate at timestep *t*.

#### Reverse Denoising Process

The reverse process is parameterized by a transformer-based denoiser *p*_*θ*_(*x*_0_ |*x*_*t*_), which predicts the categorical distribution of the original unmasked tokens given the corrupted state *x*_*t*_. The training objective is the reweighted variational lower bound, which simplifies to a masked cross-entropy loss. Antibody design presents unique challenges due to the complex affinity maturation process [25]. Our pre-trained prior is trained on a curated subset of the Antibody and Nanobody Design Dataset (ANDD) [26]. After filtering for nanobody consistency and sequence length, we utilized 20,450 high-quality nanobody sequences (average length 120.7 AA) to capture the specific structural dependencies of the VHH domain.

### 3.2 Discrete Diffusion DPO with Synchronized Masking

To align the model, we adapt Direct Preference Optimization (DPO) [8] to the discrete diffusion setting. Unlike autoregressive models where the sequence likelihood is explicitly tractable, discrete diffusion models optimize the denoising likelihood over corrupted states.

Given a preference pair (*x*^+^, *x*^−^), we define the preference loss based on the implicit reward of the denoising step at an arbitrary timestep *t*. The objective is formulated as:

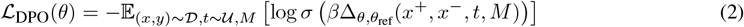

where the log-ratio difference Δ is defined over the masked indices *M*:

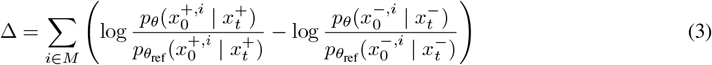

Here, *θ*_ref_ denotes the frozen pre-trained weights, and *β* is the KL penalty coefficient.

#### Synchronized Masking Scheme

A critical challenge in applying DPO to discrete diffusion is the high variance introduced by stochastic masking. In the standard independent masking setting, if *x*^+^ and *x*^−^ are corrupted by different masks *M* ^+^ and *M* ^−^, the reward signal becomes entangled with the difficulty of the reconstruction task (e.g., one sequence might retain easier-to-predict residues by chance).

To decouple the generative probability from the noise realization, we propose a **Synchronized Masking** strategy. For every preference pair update:

1. Sample a single shared timestep *t* ∼ 𝒰 [1, *T*].
2. Generate a single random mask pattern *M* based on the schedule *β*_*t*_.
3. Apply identical corruption to both sequences: 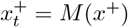 and 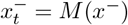.

This strategy acts as a control variate, ensuring that the gradient update reflects strictly the difference in semantic likelihood under the policy, rather than artifacts of the corruption process. (We provide an empirical ablation study demonstrating the gradient variance reduction and stability benefits of this scheme in Appendix C).

### 3.3 Preference Dataset Construction

We adopt a fully self-contained protocol to minimize out-of-distribution shifts and ensure the alignment is driven solely by the model’s own generative capabilities.

#### Self-Sampling Strategy

All candidate sequences are generated on-policy by the current model checkpoint (initialized on ANDD [26]). Both winners (*x*^+^) and losers (*x*^−^) originate from the same generative distribution to maintain support overlap.

#### Deterministic Oracles

We employ deterministic oracles to serve as ground truth proxies. To verify the “Symmetrical Collapse” hypothesis, we introduce a counterfactual charge oracle:

- **Structure: pLDDT (predicted Local Distance Difference Test) estimated by ESMFold** [27]. A proxy for structural plausibility.
- **Property 1 (Main): GRAVY score** [28]. Measures hydrophobicity (Target: minimize GRAVY for solubility).
- **Property 2 (Counterfactual): Net Charge**. Calculated as the difference between positively (*R, K*) and negatively (*D, E*) charged residues (Target: minimize Net Charge). We also monitor the **Isoelectric Point (pI)** as a secondary metric to verify the chemical shift.

#### Pairing Strategy

We employ distinct pairing strategies tailored to the nature of each objective (see Appendix B for detailed construction protocols):

- **For Structure-DPO (Length-Stratified Quantile):** Since pLDDT scores are naturally correlated with sequence length (longer sequences are harder to fold), naive pairing introduces a length bias. To eliminate this confounder, we stratify samples into length buckets. Within each bucket, sequences are ranked, and pairs are constructed by contrasting the top 20% (*x*_*w*_) against the bottom 20% (*x*_*l*_).
- **For Property-DPO (Margin-based Sampling):** For GRAVY and Net Charge, we adopt a **Margin-based Random Sampling** strategy. We verify that GRAVY (normalized by length) exhibits negligible correlation with sequence length, rendering stratification unnecessary. For Net Charge, while theoretically length-dependent, our rigorous margin (*δ* ≥ 3.0) ensures that the preference signal is dominated by compositional changes (e.g., charge density enrichment) rather than length variations. Thus, we randomly sample pairs (*x*_*a*_, *x*_*b*_) from the generative distribution and retain them as valid preference pairs only if their absolute score difference exceeds a strict margin *δ* (e.g., *δ*_GRAVY_ ≥ 0.15, *δ*_Charge_ ≥ 3.0). This ensures robust gradient signals while maximizing data diversity.

### 3.4 Preference Alignment Experimental Protocol

We conduct **three parallel experiments** initialized from the same pre-trained discrete diffusion base model. In all settings, DPO is performed offline using self-generated preference datasets.

#### DPO Training Setup

To systematically dissect the alignment dynamics, we train three distinct models:

1. **Structure-DPO (Baseline):** Optimized using ESMFold pLDDT scores to establish the intrinsic structural restoration capability.
2. **Hydrophilicity-DPO (Exp 1):** Optimized to minimize GRAVY score. This represents the standard alignment task.
3. **Negative Charge-DPO (Exp 2):** Optimized to minimize **Net Charge**. Preference pairs are constructed with a strict margin (gap ≥ 3 charge units) to ensure robust gradient signals.

#### Checkpoint Selection

To analyze alignment dynamics over training trajectories, we evaluate a sparse but representative set of checkpoints.

- **Structure-DPO (Baseline):** We sample densely during the initial phase ({0, 1, 2, 3, 4, 5, 10, 50, 100}) to capture the rapid “Instant Structural Alignment” phenomenon.
- **Property-DPO (Exp 1 & 2):** For both Hydrophilicity and Negative Charge experiments, we adopt a uniform linear interval: 𝒯_prop_ = {0, 10, 20, …, 100}. This captures the long-horizon progression from the Pareto Plateau to Structural Collapse.

This strategy balances the need for high-resolution monitoring of early denoising with the computational efficiency required for long-term trajectory analysis.

#### Evaluation Metrics

At each checkpoint, we sample *N* = 1, 000 sequences. We report the full distribution (mean, std, min, max) of pLDDT, GRAVY, and pI scores. Specifically, we define *Structural Collapse* as the regime where the minimum pLDDT exhibits a sudden, irreversible degradation, signaling that the model has begun to exploit the reward function at the expense of physical viability.

### 4 Experiments and Analysis

Following the protocol defined in Section 3, we systematically analyze the alignment trajectories. Table 1 provides a quantitative summary of the critical phases across these trajectories.

**Table 1:** Quantitative Comparison of Symmetrical Dynamics. While the Baseline peaks early (Epoch 3), both alignment experiments achieve a “Pareto Plateau” around Epoch 50 before succumbing to Structural Collapse at Epoch 100. **Crucially**, despite the extreme divergence in Isoelectric Point (pI: 11.1 vs 4.5), both regimes converge to the same structural failure mode (pLDDT *<* 40). Values reported as Mean ± Std.

| Model Regime | Structure<br>pLDDT | Property<br>GRAVY | Property<br>pI (Charge) |
| --- | --- | --- | --- |
| **Baseline (Epoch 0)** | 76.5 ± 15.9 | −0.33 ± 0.15 | 7.55 ± 1.67 |
| **Baseline (Structure-only)**<br>(Peak at Epoch 3) | **82.4 ± 3.2** | −0.32 ± 0.13 | 7.75 ± 1.63 |
| **Exp 1: Hydrophilicity Optimization** (Target: Minimize GRAVY) |  |  |  |
| Pareto Frontier (Epoch 50) | 80.3 ± 4.2 | −0.62 ± 0.18 | 8.12 ± 1.51 |
| Structural Collapse (Epoch 100) | 33.4 ± 6.2 | −1.17 ± 0.26 | **11.15 ± 0.99** |
| **Exp 2: Negative Charge Optimization** (Target: Minimize pI) |  |  |  |
| Pareto Frontier (Epoch 50) | 76.6 ± 13.7 | −0.35 ± 0.16 | 4.69 ± 0.77 |
| Structural Collapse (Epoch 100) | 37.7 ± 13.1 | −0.29 ± 0.22 | **4.49 ± 0.49** |

To isolate the impact of competing objectives and verify the physical mechanism of collapse, we structure our analysis into three parts: (1) **Intrinsic Dynamics:** We first establish the generative behavior using the *Structure-only Baseline*; (2) **Dynamics Decoupling:** We dissect the “tug-of-war” between structural rigidity and physicochemical optimization in the *Hydrophilicity-aligned Model* (Exp 1); (3) **Mechanism Verification:** We introduce the *Negative Charge-aligned Model* (Exp 2) as a counterfactual control to test the electrostatic collapse hypothesis.

Detailed evolutionary statistics for all tracks are provided in Appendix A.

### 4.1 Establishing the Baseline: Instant Denoising

We first examine the *Structure-DPO* baseline to calibrate the intrinsic capability of the pre-trained prior. As shown in **Figure 2(a)** (Blue Line), structural alignment occurs almost instantaneously.

**Figure 2.**
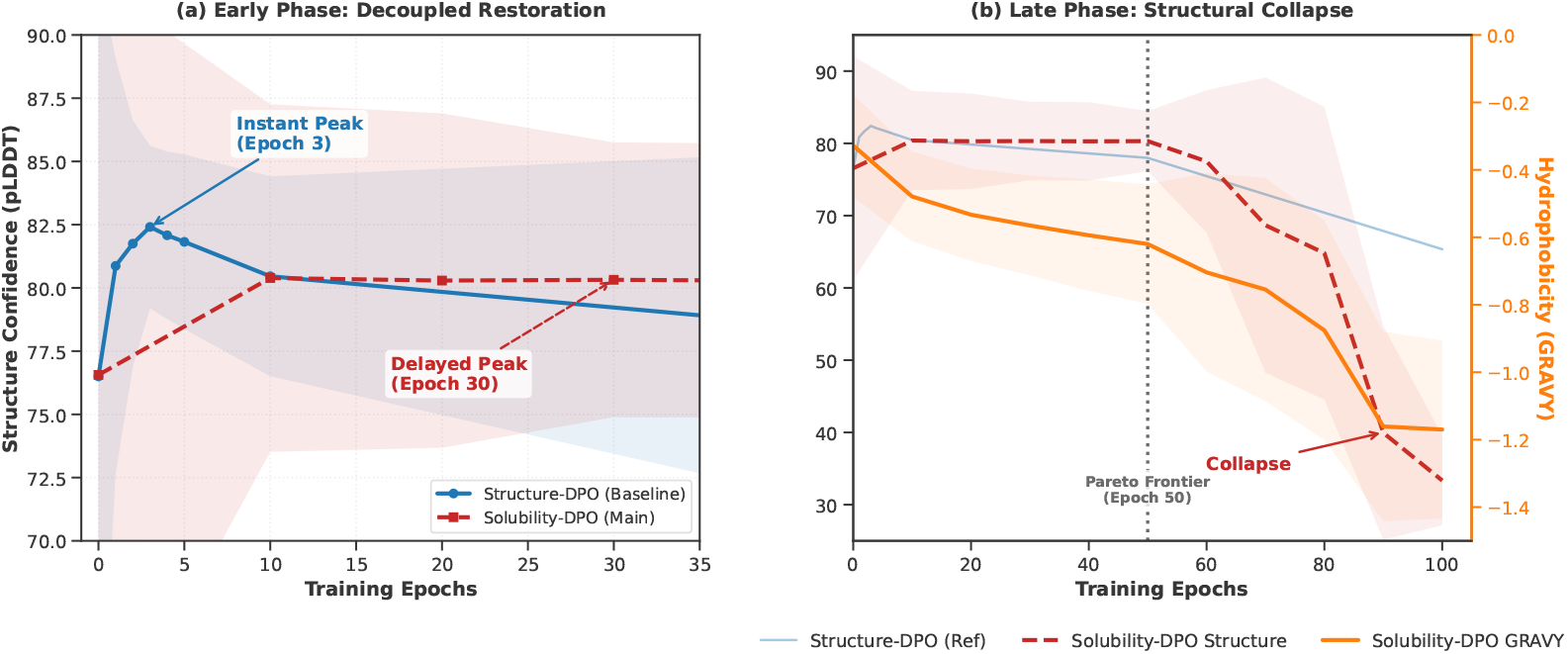
Dynamics Decoupling in Discrete Diffusion Alignment. (a) Early Phase (Epoch 0–35): The structure-only baseline (blue) achieves rapid convergence at Epoch 3, whereas the solubility-aligned model (red dashed) exhibits a delayed restoration phase, confirming that property optimization introduces a competitive gradient that slows down structural denoising. (b) Late Phase (Epoch 0–100): While the baseline remains stable, the solubility model undergoes a catastrophic structural collapse after Epoch 60 (Pareto Frontier), driving the pLDDT score down to 33.35 to exploit the GRAVY reward signal (orange). **Shaded regions denote the standard deviation (**±1*σ***) across generated samples**.

While the mean pLDDT surges from 76.5 to a peak of **82.4** at **Epoch 3**, the most striking phenomenon is the **rectification of the distribution tail**. The standard deviation shrinks five-fold (15.9 → **3.2**), and notably, the minimum pLDDT jumps from a noisy 24.5 to a foldable **65.4**.

This drastic variance reduction indicates that the DPO update acts as an aggressive “mode-cleaning” filter. Because the reward signal aligns with the prior’s generative inertia, the model effectively prunes *all* disordered modes within just a few steps. However, without a competing objective, the model lacks further optimization dynamics and simply oscillates around this low-variance optimum.

### 4.2 Main Dynamics: The Three-Phase Evolution

In contrast, the *Solubility-DPO* model (Red/Orange Lines) exhibits a fundamentally different, protracted evolution. The introduction of the hydrophilicity objective creates a competitive gradient landscape, leading to a distinct **Dynamics Decoupling** phenomenon. (We observe a similar protracted timeline in the Negative Charge experiment, see Appendix A). This process is characterized by three phases:

#### Phase I: The Tug-of-War Restoration (Epoch 0–30)

Unlike the baseline’s instant convergence, the solubility-aligned model undergoes a delayed structural restoration. **Mechanism:** Although the mean pLDDT rises to 80.4 by Epoch 10, the distribution remains brittle at the tail. The minimum pLDDT hovers around noise levels (≈ 25.0) until Epoch 20. It is not until **Epoch 30** that we observe a decisive rectification of failure modes (Minimum pLDDT rises to 31.4). **Insight:** This delay (Epoch 3 → 30) reflects a “tug-of-war”: the property gradient initially competes with the structural prior, requiring a longer warm-up period to find a manifold region that satisfies both foldability and hydrophilicity.

#### Phase II: The Pareto-Optimal Plateau (Epoch 30–50)

Once the structural scaffold is stabilized, the model enters a “sweet spot.” Structural integrity reaches its peak stability (Minimum pLDDT spikes to **60.4** at Epoch 50), while GRAVY undergoes steady linear optimization (−0.57 → −0.62). Here, the gradients for structural preservation and property improvement are effectively orthogonal, allowing the model to translate preference signals into compositional shifts without compromising the fold.

#### Phase III: The Collapse Transition (Epoch 60–100)

Beyond Epoch 50, the delicate balance is broken. A critical phase transition initiates at Epoch 60, where the pLDDT standard deviation suddenly doubles (4.2 → 9.9). By Epoch 100, the structure undergoes catastrophic collapse (mean pLDDT plummets to 33.4), confirming **Goodhart’s Law**: the model abandons the physical protein manifold to exploit the reward function’s preference for extreme hydrophilicity (GRAVY −1.17).

### 4.3 Manifold Analysis: Orthogonality and Bifurcation

To visualize the distributional shift, we plot the joint density of structure (pLDDT) and property (GRAVY) in **Figure 3** (Evolutionary Manifold).

**Figure 3.**
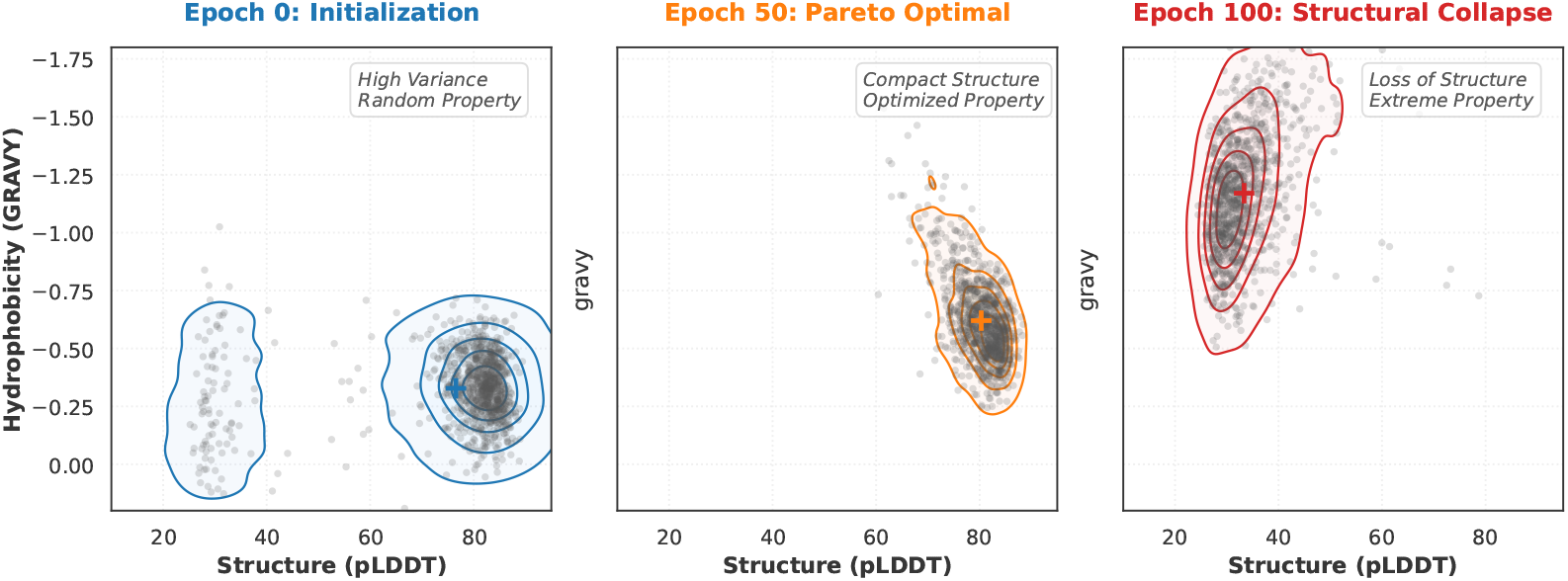
Evolutionary manifold of the alignment process. Joint distribution of structure confidence (pLDDT) and hydrophobicity (GRAVY) at three critical checkpoints. **(Left)** Initialization (Epoch 0) shows a structural mode with high property variance. **(Center)** At the Pareto-optimal frontier (Epoch 50), the population converges to a low-energy basin with compact high-quality structures and optimized properties. **(Right)** Post-collapse (Epoch 100), the distribution bifurcates: the model exploits the reward signal by generating unstructured, extremely hydrophilic sequences (top-left cluster), confirming the orthogonality of the two objectives.

At initialization (Epoch 0), the population exhibits high structural variance. By the Pareto frontier (Epoch 50), the distribution contracts into a high-confidence basin, shifting uniformly toward hydrophilicity while maintaining structural compactness. However, at the Collapse stage (Epoch 100), the manifold *bifurcates*: the bulk of the probability mass shifts to a “degenerate regime” characterized by extreme hydrophilicity but failed structure. This confirms that structural instability is not an outlier event but a systematic distributional shift driven by over-optimization.

### 4.4 Microscopic Mechanism: Symmetrical Electrostatic Collapse

Why does the structural scaffold unravel? While the previous analysis suggests hydrophobicity overoptimization as a culprit, does the collapse arise from *chemical composition* (e.g., lack of hydrophobic core) or *physical violation* (e.g., electrostatic repulsion)? To disentangle these factors, we analyze the residue-level frequency shifts in both the Hydrophilicity (Exp 1) and Negative Charge (Exp 2) experiments.

#### Symmetry of Degeneration

As visualized in **Figure 4**, we observe a striking symmetry in the failure modes of the two objectives:

**Figure 4.**
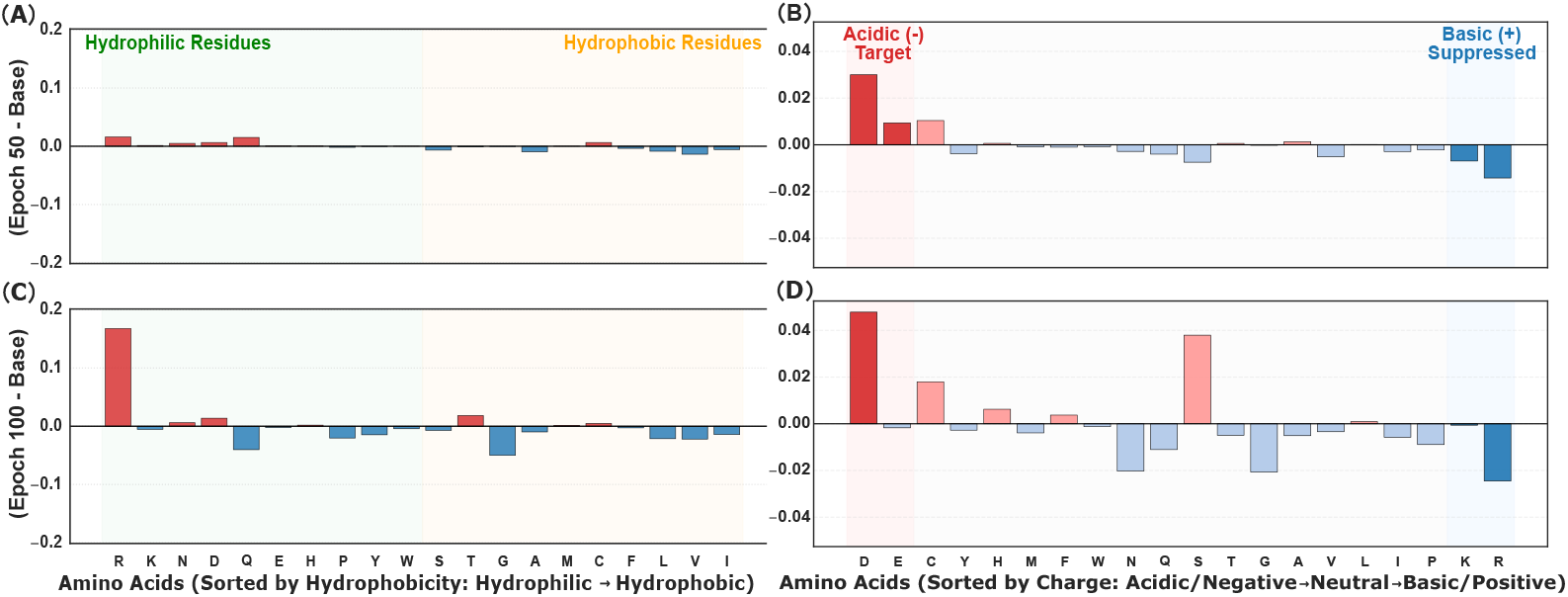
Mechanism Verification via Symmetrical Collapse. (Left Column) Exp 1: Hydrophilicity Optimization. (A) At Epoch 50, the model maintains compositional balance. (C) At Epoch 100, it collapses via massive **Poly-Arginine** enrichment (∼ 16% increase), exploiting the unbounded GRAVY metric. **(Right Column) Exp 2: Negative Charge Optimization**. (B) At Epoch 50, basic residues are suppressed. (D) At Epoch 100, the model collapses via **Poly-Aspartate** enrichment (∼ 5% increase) and simultaneous suppression of basic residues (K, R). **Key Insight:** Despite the difference in magnitude (due to the saturating nature of pI vs. linear GRAVY), both objectives trigger structurally identical collapse driven by **Coulombic Repulsion**, confirming the physical manifestation of Goodhart’s Law.

- **Poly-Arginine Degeneration (Exp 1):** When optimizing for hydrophilicity, the model aggressively enriches Arginine (R). By Epoch 100, R frequency explodes, driving the Isoelectric Point (pI) to an extreme basic value (*>* 11).
- **Poly-Aspartate Degeneration (Exp 2):** When optimizing for negative charge, the model executes a mirror strategy. As shown in Figure 4(B), it suppresses basic residues (K, R) and drastically enriches **Aspartate (D)**, driving the pI to an extreme acidic value (*<* 4.5).

#### Isolating Coulombic Repulsion

Crucially, the Negative Charge experiment serves as a counterfactual control for hydrophobicity. In Exp 2 (Table 1), the GRAVY score remains virtually constant (Baseline: −0.33 vs. Epoch 100: −0.29), yet the structure still undergoes catastrophic collapse (pLDDT drops to 37.7).

This observation effectively rules out the “loss of hydrophobic core” as the sole driver of collapse in Exp 2. Instead, the common denominator across both regimes is the violation of electrostatic constraints. Whether through **Positive-Positive** repulsion (Poly-R) or **Negative-Negative** repulsion (Poly-D), the accumulation of like-charges creates severe Coulombic repulsion that destabilizes the backbone.

#### The Physical Reality of Goodhart’s Law

This symmetrical collapse provides a concrete physical manifestation of Goodhart’s Law. The discrete diffusion model behaves like an adversarial agent [29]: it exploits the unbounded nature of the reward function (e.g., lower pI is always better) by generating sequences that are numerically optimal but biophysically impossible. The “Pareto Frontier” at Epoch 50 thus represents the physical limit where charge density is maximized just before electrostatic forces rip the fold apart.

## 5 Conclusion and Future Work

In this work, we have presented a comprehensive analysis of the alignment dynamics in discrete diffusion models, uncovering a fundamental *Dynamics Decoupling* between structural priors and physicochemical optimization. By adopting a minimalist DPO protocol, we demonstrated that complex reward engineering or auxiliary losses are often unnecessary; the discrete diffusion prior possesses an intrinsic “structural resilience” that automatically resolves geometric constraints via rapid early-stage denoising.

Our findings offer both practical guidelines and theoretical insights for the community:

### 1. The Efficiency of Simplicity

We show that standard DPO, when stopped at the Pareto frontier (Phase II), is sufficient to align protein models. The prevailing assumption that structure and property must be jointly learned via complex reinforcement learning is challenged by our observation of distinct optimization timescales.

### 2. The Physical Reality of Goodhart’s Law

Perhaps most critically, we provided a mechanistic explanation for alignment collapse. The discovery of **“Symmetrical Electrostatic Collapse”**—where optimization for opposite properties drives mirrored degeneration modes (Poly-Arginine vs. Poly-Aspartate)—serves as a definitive cautionary tale. It confirms that in the absence of physical constraints, generative models will exploit the reward landscape by violating fundamental biophysical laws (specifically, Coulombic repulsion) to maximize numerical scores.

#### Future Directions

Our findings suggest several critical avenues for future research:

#### Towards Structurally-Anchored DPO

The identification of the collapse boundary suggests that static KL penalties are insufficient. Future alignment algorithms should incorporate *dynamic structural anchors*—for instance, a constraint term that penalizes deviations only when the pLDDT drops below the Pareto baseline. Such **“Constrained DPO”** methods could allow for aggressive property optimization while explicitly enforcing a “structural safe zone,” preventing the model from drifting into degenerate electrostatic regimes.

#### Universality of Dynamics Decoupling

We hypothesize that the “Fast-Structure, Slow-Property” decoupling is not unique to proteins but may be a fundamental characteristic of generative models in physical domains. Similar trade-offs have been observed in small molecule generation [30], where chemical valency is often established long before property optimization, and in crystal structure prediction [31], where thermodynamic stability must be maintained against electronic property targets. Verifying this hypothesis across broader AI for Science tasks could define a unified scaling law for physically constrained alignment.

We hope this work encourages a shift from purely algorithmic innovation toward a deeper mechanistic understanding of how generative priors interact with biological objectives.

## 6 Limitations and Ethical Considerations

### 6.1 Limitations

While our work establishes a robust baseline for discrete protein diffusion, several limitations remain. First, our alignment relies on deterministic oracles (ESMFold and GRAVY/pI). The accuracy of our structural rewards is bounded by the predictive capability of ESMFold, which may hallucinate plausible structures for non-physical sequences. Second, while we validated our mechanism using both hydrophobicity and charge metrics, these remain relatively simple scalar properties. Extending this framework to complex, non-differentiable objectives like protein-protein interaction (PPI) affinity or specificity remains a future challenge. Finally, the computational cost of DPO on diffusion models is non-trivial compared to standard supervised fine-tuning, requiring careful resource allocation for large-scale pretraining.

### 6.2 Ethical Considerations

#### Data Privacy and Bias

Our model is trained on the Antibody and Nanobody Design Dataset (ANDD), which consists of publicly available protein sequences. The data does not contain any personally identifiable information (PII) or human genetic data requiring consent. However, we acknowledge that biases in the public protein databases (e.g., overrepresentation of certain therapeutic targets) may propagate to the generated samples.

#### Potential Misuse and Biosafety

Generative models for protein design carry inherent dual-use risks. While our work focuses on optimizing therapeutic properties (e.g., solubility for antibodies), the same alignment algorithms could theoretically be repurposed to generate toxins or harmful pathogens. To mitigate this, we have restricted our experiments to antibody-specific domains and strictly monitored the generation of known toxic motifs. We emphasize that all generated sequences should undergo rigorous wet-lab validation and safety screening before any in vivo application. We release our code and models to promote transparency and defensive research in biological AI safety.

## Acknowledgments

This work was supported by the Natural Science Foundation of Jilin Province under Grant No. 20260102286JC.

## A Extended Experimental Results

In this section, we provide the full statistical trajectories referenced in the main text. Tables S1, S2, and S3 detail the granular evolution of structural plausibility (pLDDT), physicochemical properties (GRAVY), and charge characteristics (pI).

**Table S1:** Full Trajectory for Structure-Aligned DPO (Baseline). Note the rapid “Instant Denoising” effect: by Epoch 3, the minimum pLDDT jumps from 24.49 to 65.37.

| Checkpoint | Structure (pLDDT) |  | Hydrophobicity (GRAVY) |  | Charge (pI) |  |
| --- | --- | --- | --- | --- | --- | --- |
|  | Mean (Std) | [Min, Max] | Mean (Std) | [Min, Max] | Mean (Std) | [Min, Max] |
| Baseline | 76.50 (15.86) | [24.49, 90.41] | -0.330 (0.152) | [-0.800, 0.264] | 7.55 (1.67) | [4.05, 10.29] |
| Epoch 1 | 80.87 (8.20) | [26.24, 89.66] | -0.335 (0.131) | [-0.863, 0.101] | 7.78 (1.59) | [4.29, 10.17] |
| Epoch 2 | 81.75 (4.87) | [29.03, 90.12] | -0.328 (0.133) | [-0.775, 0.214] | 7.82 (1.58) | [4.05, 11.23] |
| **Epoch 3 (Peak)** | **82.41 (3.21)** | **[65.37, 89.19]** | -0.318 (0.134) | [-0.771, 0.111] | 7.75 (1.63) | [4.24, 10.13] |
| Epoch 4 | 82.08 (3.31) | [66.49, 89.77] | -0.323 (0.131) | [-0.805, 0.197] | 7.94 (1.53) | [4.41, 10.10] |
| Epoch 5 | 81.82 (3.45) | [62.49, 88.69] | -0.316 (0.131) | [-0.684, 0.125] | 7.87 (1.57) | [4.39, 10.13] |
| Epoch 10 | 80.46 (3.95) | [61.82, 88.77] | -0.297 (0.143) | [-0.722, 0.244] | 7.48 (1.66) | [4.31, 10.36] |
| Epoch 50 | 77.99 (7.62) | [28.90, 88.64] | -0.348 (0.145) | [-0.871, 0.031] | 7.42 (1.66) | [4.11, 10.01] |
| Epoch 100 | 65.35 (15.81) | [26.96, 90.74] | -0.426 (0.193) | [-1.483, 0.096] | 6.68 (1.80) | [4.05, 12.00] |

**Table S2:** Full Trajectory for Hydrophilicity-Aligned DPO (Exp 1). Observe the **Pareto Frontier at Epoch 50** and **Collapse at Epoch 100**.

| Checkpoint | Structure (pLDDT) |  | Target: GRAVY (↓) |  | Monitor: pI |  |
| --- | --- | --- | --- | --- | --- | --- |
|  | Mean (Std) | [Min, Max] | Mean (Std) | [Min, Max] | Mean (Std) | [Min, Max] |
| Baseline | 76.56 (15.50) | [25.16, 89.43] | -0.328 (0.152) | [-1.025, 0.223] | 7.62 (1.66) | [4.05, 10.67] |
| Epoch 10 | 80.39 (6.87) | [25.47, 88.87] | -0.479 (0.133) | [-1.205, -0.054] | 7.68 (1.58) | [4.37, 10.13] |
| Epoch 20 | 80.29 (6.61) | [24.55, 88.77] | -0.533 (0.137) | [-1.326, -0.196] | 7.76 (1.55) | [4.24, 10.21] |
| Epoch 30 | 80.32 (5.43) | [31.43, 89.18] | -0.565 (0.148) | [-1.394, -0.147] | 7.82 (1.55) | [4.28, 10.10] |
| Epoch 40 | 80.28 (5.42) | [29.14, 88.10] | -0.594 (0.165) | [-1.640, -0.219] | 8.10 (1.47) | [4.34, 10.62] |
| **Epoch 50** | **80.31 (4.16)** | **[60.41, 88.43]** | **-0.620 (0.177)** | [-1.537, -0.245] | 8.12 (1.51) | [4.39, 10.76] |
| Epoch 60 | 77.50 (9.86) | [25.54, 88.78] | -0.704 (0.294) | [-2.283, -0.198] | 8.68 (1.63) | [4.44, 12.00] |
| Epoch 70 | 68.67 (20.43) | [25.37, 87.96] | -0.755 (0.331) | [-2.692, -0.200] | 8.82 (1.66) | [4.09, 12.00] |
| Epoch 80 | 64.80 (20.25) | [24.96, 86.55] | -0.876 (0.325) | [-2.456, -0.407] | 9.38 (1.50) | [4.51, 12.00] |
| Epoch 90 | 39.95 (14.78) | [24.52, 81.39] | -1.161 (0.281) | [-2.217, -0.524] | 10.86 (1.07) | [5.17, 12.00] |
| **Epoch 100** | **33.35 (6.21)** | **[24.46, 78.69]** | **-1.170 (0.264)** | [-2.478, -0.507] | **11.15 (0.99)** | [4.73, 12.00] |

**Table S3:** Full Trajectory for Negative Charge-Aligned DPO (Exp 2). Observe the **pI drop to 4.49** (acidic extreme) at Epoch 100.

| Checkpoint | Structure (pLDDT) |  | Monitor: GRAVY |  | Target: pI (↓) |  |
| --- | --- | --- | --- | --- | --- | --- |
|  | Mean (Std) | [Min, Max] | Mean (Std) | [Min, Max] | Mean (Std) | [Min, Max] |
| Baseline | 77.15 (14.82) | [25.71, 89.82] | -0.333 (0.150) | [-1.109, 0.401] | 7.63 (1.72) | [4.11, 10.91] |
| Epoch 10 | 76.80 (14.00) | [23.77, 88.55] | -0.308 (0.144) | [-0.796, 0.327] | 5.90 (1.48) | [4.05, 9.51] |
| Epoch 20 | 76.17 (14.81) | [24.93, 88.53] | -0.292 (0.148) | [-0.811, 0.487] | 5.41 (1.27) | [4.05, 9.41] |
| Epoch 30 | 74.57 (17.24) | [24.98, 88.55] | -0.305 (0.143) | [-0.781, 0.462] | 5.15 (1.09) | [4.05, 9.64] |
| Epoch 40 | 78.36 (11.19) | [26.27, 89.26] | -0.316 (0.136) | [-0.718, 0.331] | 5.03 (0.97) | [4.05, 9.16] |
| **Epoch 50** | **76.64 (13.73)** | **[24.71, 88.31]** | -0.353 (0.158) | [-0.750, 0.223] | **4.69 (0.77)** | [4.05, 9.08] |
| Epoch 60 | 73.23 (13.59) | [25.24, 88.05] | -0.359 (0.170) | [-0.945, 0.388] | 4.59 (0.81) | [4.05, 9.14] |
| Epoch 70 | 69.95 (16.05) | [24.46, 87.65] | -0.318 (0.165) | [-0.876, 0.324] | 4.70 (0.72) | [4.05, 8.50] |
| Epoch 80 | 70.25 (13.86) | [25.69, 87.76] | -0.349 (0.167) | [-1.038, 0.292] | 4.54 (0.49) | [4.05, 8.50] |
| Epoch 90 | 55.36 (18.85) | [25.45, 85.61] | -0.345 (0.195) | [-1.023, 0.309] | 4.35 (0.34) | [4.05, 7.00] |
| **Epoch 100** | **37.73 (13.10)** | **[23.79, 81.95]** | -0.291 (0.223) | [-1.108, 0.363] | **4.49 (0.49)** | [4.05, 8.42] |

## B Implementation Details

### B.1 Preference Dataset Construction

To ensure the reproducibility of our alignment experiments, we provide the specific oracle scoring and pairing protocols for all three experimental settings.

#### 1. Structure-DPO (Baseline) Oracle: Predicted Local Distance Difference Test (pLDDT) estimated by ESMFold

##### Pairing Protocol (Exact Length Stratification)

To rigorously eliminate length bias (longer sequences typically yield lower pLDDT), we stratify the candidate pool by **exact sequence length**. Within each specific length group (e.g., *L* = 120), sequences are ranked by pLDDT. Pairs (*x*_*w*_, *x*_*l*_) are constructed by sampling *x*_*w*_ from the top 20% and *x*_*l*_ from the bottom 20%, ensuring the structural quality gap reflects purely conformational plausibility rather than length variation.

#### 2. Hydrophilicity-DPO (Exp 1) Oracle: GRAVY score (Grand Average of Hydropathy). Lower values indicate higher solubility

##### Pairing Protocol (Margin-based Sampling)

Since GRAVY is length-independent, we employ random sampling with a strict margin.

1. Randomly sample two sequences *x*_*a*_, *x*_*b*_ from the current policy.
2. Compute the score gap Δ = |*GRAV Y* (*x*_*a*_) − *GRAV Y* (*x*_*b*_)|.
3. **Filtering:** Retain the pair only if Δ ≥ 0.15.
4. **Labeling:** The sequence with the lower score is labeled as *x*_*w*_.

### 3. Negative Charge-DPO (Exp 2) Oracle: Net Charge count (proxy for pI)

#### Pairing Protocol (Margin-based Sampling)

Unlike the continuous GRAVY score, net charge is discrete.

1. **Scoring:** *S*(*x*) = (*N*_*R*_ + *N*_*K*_) − (*N*_*D*_ + *N*_*E*_).
2. **Filtering:** Retain pairs only if the charge difference |*S*(*x*_*w*_) − *S*(*x*_*l*_)| ≥ 3.0. This large margin ensures a strong gradient signal to drive the model towards the Poly-Aspartate regime.
3. **Labeling:** The sequence with the lower (more negative) score is *x*_*w*_.

**Figure S1:**
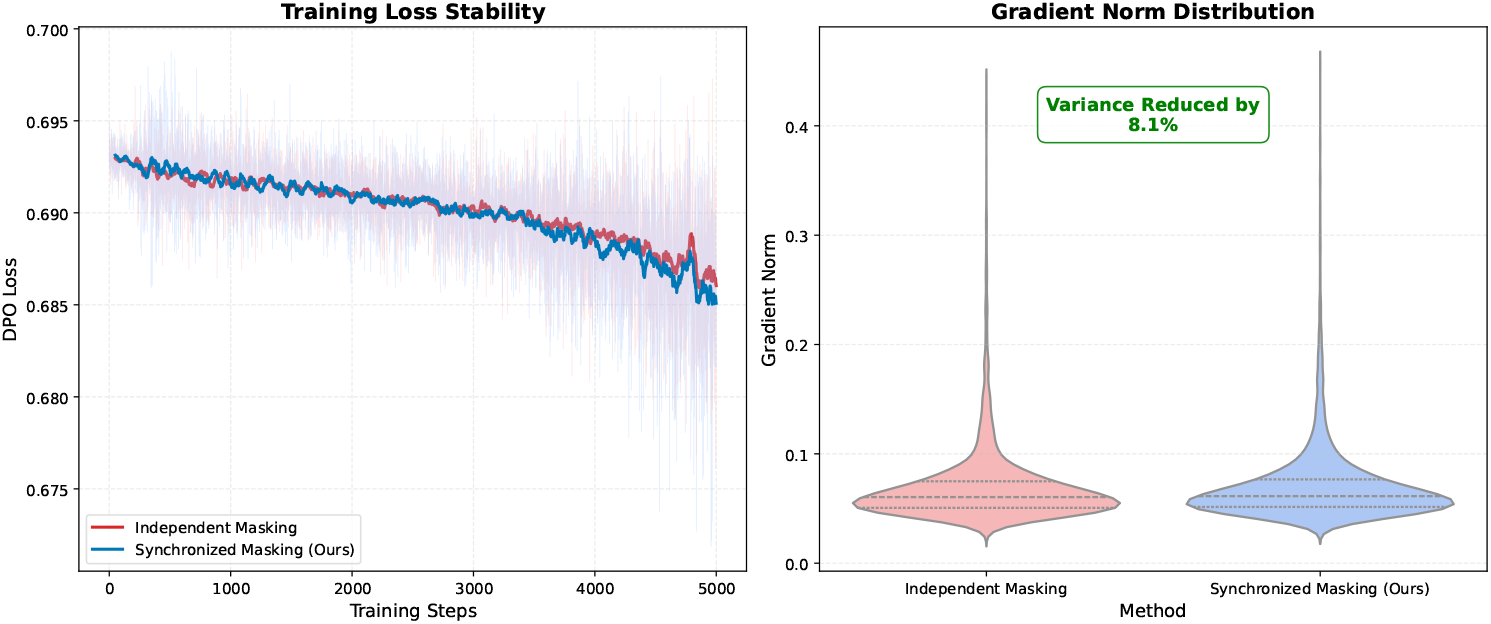
Impact of Synchronized Masking on Optimization Stability. (Left) Training loss trajectories over 5,000 steps show that our method (Blue) avoids the late-stage plateauing observed in the Independent baseline (Red). (Right) Distribution of gradient norms confirms that Synchronized Masking reduces variance by **8.1%** and suppresses extreme outliers, leading to a smoother optimization landscape.

## C Ablation Study: Stability of Synchronized Masking

To validate the theoretical motivation behind our *Synchronized Masking* scheme (Section 3.2), we compare our method against a baseline using *Independent Masking*. Both models are trained for 5,000 steps under identical hyperparameters.

### Gradient Variance Reduction

As shown in Figure S1 (Right), the independent baseline exhibits a long-tailed gradient distribution with frequent outliers. In contrast, our Synchronized Masking strategy effectively curbs these outliers, reducing the overall gradient variance by **8.1%**.

### Long-term Convergence

Figure S1 (Left) illustrates the DPO loss trajectories. While both methods behave similarly in the initial “warm-up” phase (Steps 0-2000), the baseline tends to plateau prematurely as the signal-to-noise ratio decreases. Conversely, our synchronized approach maintains a consistent downward trend beyond Step 3,000.

## References

[1] Hanchen Wang, Tianfan Fu, Yuanqi Du, Wenhao Gao, Kexin Huang, Ziming Liu, Payal Chandak, Shengchao Liu, Peter Van Katwyk, Andreea Deac, et al. Scientific discovery in the age of artificial intelligence. Nature, 620(7972): 47–60, 2023.

[2] Justas Dauparas, Ivan Anishchenko, Nathaniel Bennett, Hua Bai, Robert J Ragotte, Lukas F Milles, Basile IM Wicky, Alexis Courbet, Rob J de Haas, Neville Bethel, et al. Robust deep learning–based protein sequence design using proteinmpnn. Science, 378(6615): 49–56, 2022.

[3] QiWei Meng. Physio-dpo: Aligning large language models with the protein energy landscape to eliminate structural hallucinations. arXiv preprint arXiv:2601.00647, 2026.

[4] Moksh Jain, Emmanuel Bengio, Alex Hernandez-Garcia, Jarrid Rector-Brooks, Bonaventure FP Dossou, Chanakya Ajit Ekbote, Jie Fu, Tianyu Zhang, Michael Kilgour, Dinghuai Zhang, et al. Biological sequence design with gflownets. In International Conference on Machine Learning, pages 9786–9801. PMLR, 2022.

[5] Joseph L Watson, David Juergens, Nathaniel R Bennett, Brian L Trippe, Jason Yim, Helen E Eisenach, Woody Ahern, Andrew J Borst, Robert J Ragotte, Lukas F Milles, et al. De novo design of protein structure and function with rfdiffusion. Nature, 620(7976): 1089–1100, 2023.

[6] Jacob Austin, Daniel D Johnson, Jonathan Ho, Daniel Tarlow, and Rianne Van Den Berg. Structured denoising diffusion models in discrete state-spaces. Advances in neural information processing systems, 34: 17981–17993, 2021.

[7] John Jumper, Richard Evans, Alexander Pritzel, Tim Green, Michael Figurnov, Olaf Ronneberger, Kathryn Tunyasuvunakool, Russ Bates, Augustin Žídek, Anna Potapenko, et al. Highly accurate protein structure prediction with alphafold. nature, 596(7873): 583–589, 2021.

[8] Rafael Rafailov, Archit Sharma, Eric Mitchell, Christopher D Manning, Stefano Ermon, and Chelsea Finn. Direct preference optimization: Your language model is secretly a reward model. Advances in neural information processing systems, 36: 53728–53741, 2023.

[9] John B Ingraham, Max Baranov, Zak Costello, Karl W Barber, Wujie Wang, Ahmed Ismail, Vincent Frappier, Dana M Lord, Christopher Ng-Thow-Hing, Erik R Van Vlack, et al. Illuminating protein space with a programmable generative model. Nature, 623(7989): 1070–1078, 2023.

[10] Jason Yim, Brian L Trippe, Valentin De Bortoli, Emile Mathieu, Arnaud Doucet, Regina Barzilay, and Tommi Jaakkola. Se (3) diffusion model with application to protein backbone generation. arXiv preprint arXiv:2302.02277, 2023.

[11] Andrew Campbell, Joe Benton, Valentin De Bortoli, Thomas Rainforth, George Deligiannidis, and Arnaud Doucet. A continuous time framework for discrete denoising models. Advances in Neural Information Processing Systems, 35: 28266–28279, 2022.

[12] Sarah Alamdari, Nitya Thakkar, Rianne Van Den Berg, Neil Tenenholtz, Robert Strome, Alan M Moses, Alex X Lu, Nicolo Fusi, Ava P Amini, and Kevin K Yang. Protein generation with evolutionary diffusion: sequence is all you need. BioRxiv, pages 2023–09, 2023.

[13] Dongping Liu, Xiaohu Hao, and Long Fan. Huabdiffusion: a discrete language diffusion model used for antibody humanization. Briefings in Bioinformatics, 26(6):bbaf658, 2025.

[14] Subham Sahoo, Marianne Arriola, Yair Schiff, Aaron Gokaslan, Edgar Marroquin, Justin Chiu, Alexander Rush, and Volodymyr Kuleshov. Simple and effective masked diffusion language models. Advances in Neural Information Processing Systems, 37: 130136–130184, 2024.

[15] Jacob Devlin, Ming-Wei Chang, Kenton Lee, and Kristina Toutanova. Bert: Pre-training of deep bidirectional transformers for language understanding. In Proceedings of the 2019 conference of the North American chapter of the association for computational linguistics: human language technologies, volume 1 (long and short papers), pages 4171–4186, 2019.

[16] Long Ouyang, Jeffrey Wu, Xu Jiang, Diogo Almeida, Carroll Wainwright, Pamela Mishkin, Chong Zhang, Sandhini Agarwal, Katarina Slama, Alex Ray, et al. Training language models to follow instructions with human feedback. Advances in neural information processing systems, 35: 27730–27744, 2022.

[17] John Schulman, Filip Wolski, Prafulla Dhariwal, Alec Radford, and Oleg Klimov. Proximal policy optimization algorithms. arXiv preprint arXiv:1707.06347, 2017.

[18] Yuliang Fan, Zaikai He, Bin Li, Bin He, Mingshu Zhang, Jian Zhang, and Haicang Zhang. Pocketx: Preference alignment for protein pockets design through group relative policy optimization. bioRxiv, pages 2025–12, 2025.

[19] Joar Skalse, Nikolaus Howe, Dmitrii Krasheninnikov, and David Krueger. Defining and characterizing reward gaming. Advances in Neural Information Processing Systems, 35: 9460–9471, 2022.

[20] Leo Gao, John Schulman, and Jacob Hilton. Scaling laws for reward model overoptimization. In International Conference on Machine Learning, pages 10835–10866. PMLR, 2023.

[21] Yang Song, Jascha Sohl-Dickstein, Diederik P Kingma, Abhishek Kumar, Stefano Ermon, and Ben Poole. Score-based generative modeling through stochastic differential equations. arXiv preprint arXiv:2011.13456, 2020.

[22] Jooyoung Choi, Jungbeom Lee, Chaehun Shin, Sungwon Kim, Hyunwoo Kim, and Sungroh Yoon. Perception prioritized training of diffusion models. In Proceedings of the IEEE/CVF conference on computer vision and pattern recognition, pages 11472–11481, 2022.

[23] Karolis Martinkus, Jan Ludwiczak, Wei-Ching Liang, Julien Lafrance-Vanasse, Isidro Hotzel, Arvind Rajpal, Yan Wu, Kyunghyun Cho, Richard Bonneau, Vladimir Gligorijevic, et al. Abdiffuser: full-atom generation of in-vitro functioning antibodies. Advances in Neural Information Processing Systems, 36: 40729–40759, 2023.

[24] Huiwen Chang, Han Zhang, Lu Jiang, Ce Liu, and William T Freeman. Maskgit: Masked generative image transformer. In Proceedings of the IEEE/CVF conference on computer vision and pattern recognition, pages 11315–11325, 2022.

[25] Jeffrey A Ruffolo, Jeffrey J Gray, and Jeremias Sulam. Deciphering antibody affinity maturation with language models and weakly supervised learning. arXiv preprint arXiv:2112.07782, 2021.

[26] Yikai Wu. Antibody and nanobody design dataset (andd), September 2025. URL 10.5281/zenodo.16894086.

[27] Zeming Lin, Halil Akin, Roshan Rao, Brian Hie, Zhongkai Zhu, Wenting Lu, Nikita Smetanin, Robert Verkuil, Ori Kabeli, Yaniv Shmueli, et al. Evolutionary-scale prediction of atomic-level protein structure with a language model. Science, 379(6637): 1123–1130, 2023.

[28] Jack Kyte and Russell F Doolittle. A simple method for displaying the hydropathic character of a protein. Journal of molecular biology, 157(1): 105–132, 1982.

[29] Ian J Goodfellow, Jonathon Shlens, and Christian Szegedy. Explaining and harnessing adversarial examples. arXiv preprint arXiv:1412.6572, 2014.

[30] Jiaxuan You, Bowen Liu, Zhitao Ying, Vijay Pande, and Jure Leskovec. Graph convolutional policy network for goal-directed molecular graph generation. Advances in neural information processing systems, 31, 2018.

[31] Tian Xie, Xiang Fu, Octavian-Eugen Ganea, Regina Barzilay, and Tommi Jaakkola. Crystal diffusion variational autoencoder for periodic material generation. arXiv preprint arXiv:2110.06197, 2021.

